# Widespread alterations in microRNA biogenesis in human Huntington’s disease putamen

**DOI:** 10.1101/2022.01.27.478047

**Authors:** Serena Petry, Rémi Keraudren, Behnaz Nateghi, Andréanne Loiselle, Karolina Pircs, Johan Jakobsson, Chantelle Sephton, Mélanie Langlois, Isabelle St-Amour, Sébastien S. Hébert

## Abstract

Altered microRNA (miRNA) expression is a common feature of Huntington’s disease (HD) and could participate in disease onset and progression. However, little is known about the underlying causes of miRNA disruption in HD. We and others have previously shown that mutant Huntingtin (mHTT) binds to Ago2, a central component of miRNA biogenesis, and disrupts mature miRNA levels. In this study, we sought to determine if miRNA maturation *per se* was compromised in HD. Towards this end, we characterized t major miRNA biogenesis pathway components and miRNA maturation products (pri-miRNA, pre-miRNA, and mature) in human HD (N=41, Vonsattel grades HD2-4) and healthy control (N=25) subjects. Notably, the striatum (putamen) and cortex (BA39) from the same individuals were analyzed in parallel. We show that Ago2, Drosha, and Dicer were strongly downregulated in human HD at early stages of disease. Using a panel of HD-related miRNAs (miR-10b, miR-196b, miR-132, miR-212, miR-127, miR-128), we uncovered various types of maturation defects in HD brain, the most prominent occurring at the pre-miRNA to mature miRNA maturation step. Consistent with earlier findings, we provide evidence that alterations in autophagy could participate in miRNA maturation defects. Notably, most changes occurred in the striatum, which is more prone to HTT aggregation and neurodegeneration. Likewise, we observed no significant alterations in miRNA biogenesis in human HD cortex and blood, strengthening tissue-specific effects. Overall, these data provide important clues into the underlying mechanisms behind miRNA alterations in HD-susceptible tissues. Further investigations are now required to understand the biological, diagnostic, and therapeutic implications of miRNA/RNAi biogenesis defects in HD and related neurodegenerative disorders.

## Introduction

Huntington’s disease (HD) is an incurable, hereditary neurodegenerative disorder caused by a CAG trinucleotide repeat expansion in exon 1 of the Huntingtin (Htt) gene. At the protein level, this results in the generation of abnormal polyglutamine (PolyQ) repeats at the N-terminus of Htt. HD typically manifests itself in midlife with motor and cognitive symptoms associated with neurodegeneration in the striatum and to a lesser degree cortex. The molecular mechanisms leading to Htt-mediated neurodegeneration are still unresolved, although it is well recognized that abnormal regulation of gene expression is an early and key feature of HD neuropathology [21, 30, 33].

The small non-coding microRNAs (miRNAs) play a central role in gene expression regulation by promoting messenger RNA (mRNA) translation inhibition and/or degradation [41, 42]. MiRNA function is inherently related to its maturation that follows two major processing steps [12, 27]. First, the long primary miRNA transcript (pri-miRNA) is cleaved by the Drosha/DGCR8 complex to generate a ~70 nucleotide (nt) precursor miRNA (pre-miRNA). The pre-miRNA is transported to the cytoplasm by Exportin 5 where it is then cleaved by Dicer to generate a ~22 nt mature miRNA. The mature miRNA is finally loaded with Ago2 and associated proteins (e.g., TRBP) into the endogenous RNA-induced silencing complex (RISC) that binds to the 3’untranslated region (3’UTR) of target mRNAs. Interestingly, miRNAs can control diverse biological pathways by modulating one or several key target genes simultaneously [37]. Therefore, any disruption in this pathway could have deleterious consequences on gene expression networks and cell homeostasis.

Indeed, it is now well established that miRNAs are essential to the survival of striatal and cortical neurons [5, 13]. Loss of neuronal Dicer in adult mice leads to alterations in transcription, reduced brain size, behavioural defects, and decreased lifespan [6, 9], reminiscent of some features of HD. In HD mice (YAC128 model), Lee *et al.* observed a global upregulation or downregulation of mature miRNAs in early and late stages of disease, respectively [23]. These changes coincided with transient changes in Dicer, Drosha and Exportin mRNA levels. Recently, we and others have shown that mHtt binds to Ago2 protein [29, 35, 36], whereas transient overexpression of mHtt in cells and mice leads to higher Ago2 expression and widespread alterations in mature miRNA levels [29]. Furthermore, post-mortem studies have detected changes in mature miRNA expression profiles in the brains of HD mice and humans [16-18, 22, 24, 25, 28].

Despite these observations, there is surprisingly no clear evidence that miRNA maturation *per se* is defective in HD, especially in humans. This could have important implications in understanding miRNA regulation and function within cell survival pathways and current therapeutic efforts using the endogenous RISC (composed of Ago2 and Dicer) to silence mHtt [1, 8]. Towards this end, we have analyzed, for the first time, all major miRNA pathway components and maturation products (pri-miRNA, pre-miRNA, mature) in human HD tissues samples. Notably, our experiments were conducted in different tissues that were collected from patients at different stages of disease. In sum, our data implicate widespread defects in the pre-miRNA to mature miRNA step in HD, which overlaps with mHtt pathology and overt neurodegeneration in the striatum.

## Materials and Methods

### Human brain samples

Dissected frozen human putamen and matching cortical (BA39 region) tissues (0,5-1,2 g per sample) were obtained from the Harvard Brain Tissues Resource Center via NIH Neurobiobank (see **Suppl. Table 1**) as before [40]. This specific study included brain tissues from 25 control and 41 HD individuals. Frozen post-mortem tissues were prepared as described before and used for protein and RNA analysis [40]. CAG-repeat length was determined by the CHU de Québec Sequencing and Genotyping platform using a 6-FAM fluorescent primer (applied Biosystems Inc, Foster city, CA, USA) in a polymerase chain reaction (PCR) followed by determination of the product size by capillary electrophoresis in a 3130xl Genetic analyzers. We used the disease burden score (DBS) to estimate the lifetime exposure to mutant huntingtin in individuals with HD with the following equation: DBS = age at death x (CAG-repeat length - 35,5).

### Protein and RNA Extraction

Total proteins were extracted as previously described [40]. In brief, frozen tissues were mechanically homogenized in seven volumes of lysis buffer (150 nM NaCl, 50 mM Tris, 0.5% deoxycholate, 1% Triton X-100, 0.5% sodium dodecyl sulfate (SDS), complete protease inhibitor cocktail, 1 mM of sodium fluoride and 1 mM of activated orthovanadate as phosphatase inhibitor), then sonicated three times for 5 X 1-s pulses. The solution was spun at 100,000g for 20 min at 4°C. The supernatant (soluble proteins) was kept at −80°C until processed. The pellet was further homogenized in formic acid (FA) and spun for 20 min at 17,500g at 4°C. FA-soluble proteins (FA fraction) were dried before being sonicated in NuPAGE^®^ LDS sample buffer (Life Technologies) supplemented with 0.1M of dithiothreitol, incubated 10 min at 70°C and kept at −80°C until processed. Soluble proteins were quantified with Pierce™ BCA Protein Assay Kit (ThermoFisher Scientific) and mixed to the NuPAGE^®^ LDS sample buffer with 5% final volume of β-mercaptoethanol, then boiled 10 min at 95°C for Western blot analysis. Total RNA was extracted from frozen tissues using TRIzol reagent (Ambion by Life Technologies) according to the manufacturer’s instructions. Total RNA pellet was suspended in RNase free water, quantified, and diluted to a final concentration of 100 ng/μL. RNA was kept at −80°C until processed for qRT-PCR analysis.

### Western Blotting

Five to twenty micrograms of soluble proteins were separated by two different systems: 10% SDS-polyacrylamide gels (SDS-Page) and gradient 3-15% tris-acetate polyacrylamide gels for higher and lower molecular weight proteins. For the 10% SDS-Page, proteins were transferred onto a 0.45 μm nitrocellulose membrane (Bio-Rad, catalog n° 1620115) for 1 hour at RT at 100V. For the gradient gels, proteins were transferred onto a 0.45 μm methanol-activated PVDF membrane (Immobilon, Millipore) overnight at 4°C at 25V and 45 min at 4°C at 75V the next day. The membrane was blocked with 5% non-fat milk and 1% bovine serum albumin, then incubated overnight at 4°C with the appropriate primary antibodies (see **Suppl. Table 3)**. On the second day, membranes were incubated with appropriate secondary anti-IgG-HRP antibodies (Jackson ImmunoResearch: anti-mouse, catalog n°115-035-146 or anti-rabbit, catalog n°111-035-144) at RT for 1 h. The immune-reactive bands were acquired using Immobilon Western Chemiluminescent HRP Substrate (Millipore) and visualized with the Fusion FX (Vilber Lourmat, Eberhardzell, Germany) imaging system. Normalization was performed on total proteins obtained with Ponceau Red or StainFree staining. Band intensities were quantified using the ImageJ software.

### Dot blot

Two microliters of each sample were slowly spotted on nitrocellulose membrane. After drying the membrane, non-specific sites were blocked, and membrane was processed as described in the Western Blotting section. Dot intensity was normalized on the total amount of tissue used for the extraction.

### Primary microRNA Real Time Quantitative RT-PCR

The reverse transcription was performed with 500 ng of total RNA using the High-capacity cDNA reverse transcription kit (ThermoFisher Scientific, catalog n°4368814) according to the manufacturer’s instructions. Program: 25°C for 10 min, 37°C for 120 min and 85°C for 5 min. cDNA was stored at −20°C until further processing. The real-time quantitative PCR (qRT-PCR) was performed with TaqMan Fast Advanced Master mix (ThermoFisher Scientific, catalog n°4444963) according to manufacturer’s instructions. Primers were purchased from ThermoFisher Scientific (Hs03302879_pri, hsa-mir-10b; Hs03303255_pri, hsa-mir-127; Hs03303101_pri, hsa-mir-128-1; Hs03303111_pri, hsa-mir-132; Hs03293754_pri, hsa-mir-196b; Hs03302957_pri, hsa-mir-212). Primary microRNAs were normalized to the geographic mean of GAPDH and RPL32. The relative amount of each primary microRNA was calculated using the comparative Ct (2^−ΔΔ*Ct*^) method as before [39].

### Precursor microRNA Real Time Quantitative RT-PCR

The reverse transcription was performed with 500 ng of total RNA using the miScript RT II kit (Qiagen) according to the manufacturer’s instructions. The RT-PCR was performed with the Hiflex buffer, as recommended by the manufacturer to study precursor microRNAs. Program: 38°C for 60 min and 95°C for 5 min. cDNA was stored at −20°C until further processing. The qRT-PCR was performed with QuantiTect SYBR Green PCR Master Mix (Qiagen) according to the manufacturer’s instructions. miScript precursor assay primers were purchased from Qiagen (mir-10b ID: MP00003983; mir-127-1 ID: MP00004123; mir-128-1 ID: MP00004137; mir-132 MP00004179; mir-196b ID: MP00004935; mir-212 ID: MP00004256). Precursor microRNAs were normalized to SNORD95 (ID: MS00033726). The relative amount of each precursor microRNA was calculated using the comparative Ct (2^−ΔΔ*Ct*^) method.

### Mature microRNA Real Time Quantitative RT-PCR

The reverse transcription was performed with 10 ng of total RNA using the TaqMan MicroRNA Reverse transcription kit (ThermoFisher) according to the manufacturer’s instructions. Program: 16°C for 30 min, 42°C for 30 min and 85°C for 5 min. cDNA was stored at −20°C until further processing. The qRT-PCR was performed with TaqMan Fast Advanced Master mix (ThermoFisher Scientific) according to manufacturer’s instructions. miRNA assay primers were purchased from ThermoFisher Scientific (hsa-miR10b, 002218; hsa-miR127, 000452; hsa-miR128, 002216; hsa-miR132, 000457; hsa-miR196b, 002215; hsa-miR212, 000515). Mature microRNAs were normalized to the geographic mean of RNU48 (hsa-RNU48, 001006) and Let-7f (hsa-Let-7f, 000382). The relative amount of each mature microRNA was calculated using the comparative Ct (2^−ΔΔ*Ct*^) method.

### Statistical Analysis

All graphics and statistical analyses were performed using GraphPad Prism 9 Software (Graph Pad Software, Inc., La Jolla, CA, USA). Normality and lognormality tests were performed and parametric or non-parametric tests were used accordingly. When samples distribution passed the normality test, a parametric one-way analysis of variance (ANOVA) test followed by Dunnett’s multiple comparisons and parametric unpaired student t-test were performed. When samples distribution did not pass normality test, a non-parametric Kruskal-Wallis test followed by Dunn’s multiple comparisons and a non-parametric Mann Whitney student t-test were performed. The threshold for statistical significance was set to *p*-values < 0.05.

## Results

### Comparative biochemical analysis of Htt pathology in cortex and striatum

In an attempt to understand the impact of endogenous human Htt on miRNA maturation, we first evaluated Htt expression and pathology in two different brains regions affected in human HD (**see Suppl. Table 1**). We quantified the amount of Htt protein in 41 HD patients (N=10 HD2, N=23 HD3, N=8 HD4) and 25 Controls from matching striatal and cortical tissues. Consistent with earlier findings, we observed a decrease in soluble total (full-length) Htt in HD striatum using 1HU-4C8 and CH00146 antibodies (**Fig. 1A-E**). However, no significant changes in total Htt were shown in HD cortex. Using the 1HU-4C8 clone, we detected an increase in N-terminal fragments (MW ~ 40-50 KDa) in both brain regions. As expected, an increase in formic acid (FA)-insoluble aggregated mHtt was also observed in both regions using an anti-PolyQ antibody, although higher levels were apparent in the striatum (**Fig. 1H-I**). Along with these results, significant decreases in NeuN (neuronal marker), DARPP-32 (striatal neuron marker), and PSD-95 (post-synaptic marker) protein levels were observed in HD striatum while the cortex was mostly spared (**Fig. 1L-N**). Our results support that the relative expression level of these proteins is modulated between brain regions (**Suppl. Fig 1).** These data suggest that Htt aggregation is not the consequence of an increase expression level of endogenous Htt. Thus overall, in line with previous results suggesting that HD pathology starts in the striatum, the striatal tissue samples analyzed herein presented severe signs of Htt pathology and neurodegeneration when compared to the cortex of the same individuals.

**Figure 1.**
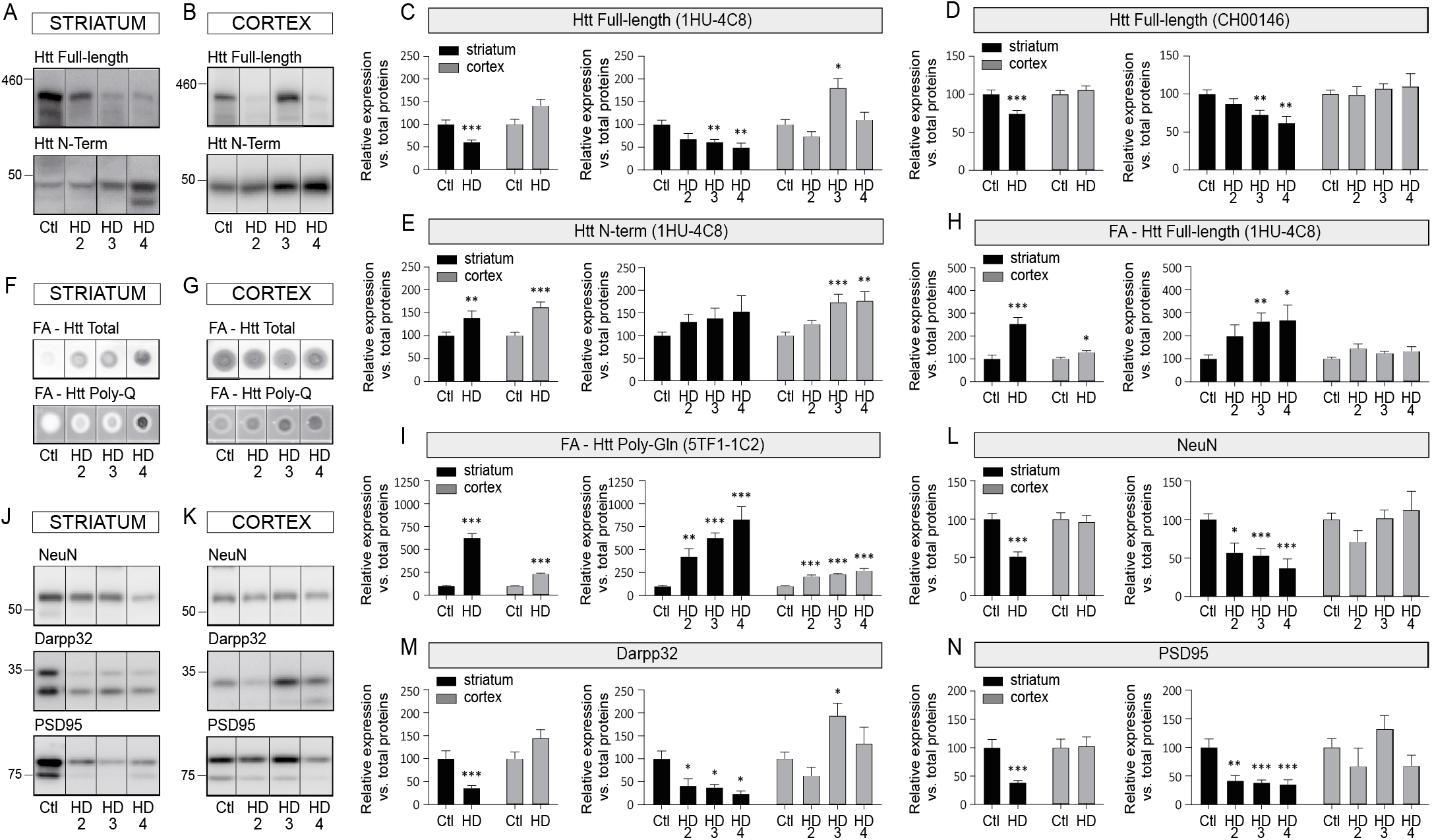
Differential Htt pathology between HD striatum and cortex. Representative immunoblots of endogenous full-length Huntingtin (Htt) (1HU-4C8 antibody) and N-terminal Htt fragments (1HU-4C8 antibody) in the soluble fraction of **(A)** the striatum or **(B)** the cortex of HD patients and Controls. **(C-E)** Protein quantifications of soluble full-length Htt (1HU-4C8 and CH00146 antibodies) and N-terminal Htt (1HU-4C8 antibody). Representative dot blots of formic acid (FA)-insoluble full-length Htt (1HU-4C8) and mutant Htt (Poly-Gln) in (**F**) the striatum and (**G**) the cortex of HD patients and Controls. (**H-I**) Protein quantifications of dot blots. Representative immunoblots of endogenous NeuN, Darpp32 and PSD95 in **(J)**the striatum or **(K)**the cortex on HD patients and Controls with quantifications in (**L-M**). Bar graphs with standard error of the mean (SEM) are shown, where the average of Controls is set as 100%. In all cases, the HD group is presented as pooled or per stage. Statistics: Ctl vs. HD as a group was calculated using a Mann-Whitney test. Ctl vs. HD stages was calculated using an analysis of covariance followed by Kruskal-Wallis multiple comparison test. Significant fold changes are provided for each group. * *P*<0.05; ** *P*<0.01; *** *P*<0.001; **** *P*<0.0001. Abbreviations: Ctl, Controls; HD, Huntington’s disease; HD2, Vonsattel grade 2; HD3, Vonsattel grade 3; HD4, Vonsattel grade 4.

### Early-stage alterations of miRNA pathway components in human HD striatum

Previous studies in mice [23, 29] showed that specific members of the miRNA biogenesis pathway are compromised in HD models. In human brains, we observed a robust decrease in Drosha, Dicer, and Ago2 protein levels from HD2 in the striatum but not in the cortex **(Fig. 2A-H)**. No significant changes in Dicer mRNA were noted in either region (**Suppl. Fig. 2**), suggesting that alterations in expression occurred at the post-transcriptional level. Modest or transient variations in DGCR8 and TRBP proteins were seen in these samples with no changes in Exportin 5. Taken together, these results suggest that core miRNA biogenesis pathway components are rapidly and specifically compromised in human HD striatum and precede overt neurodegeneration.

**Figure 2.**
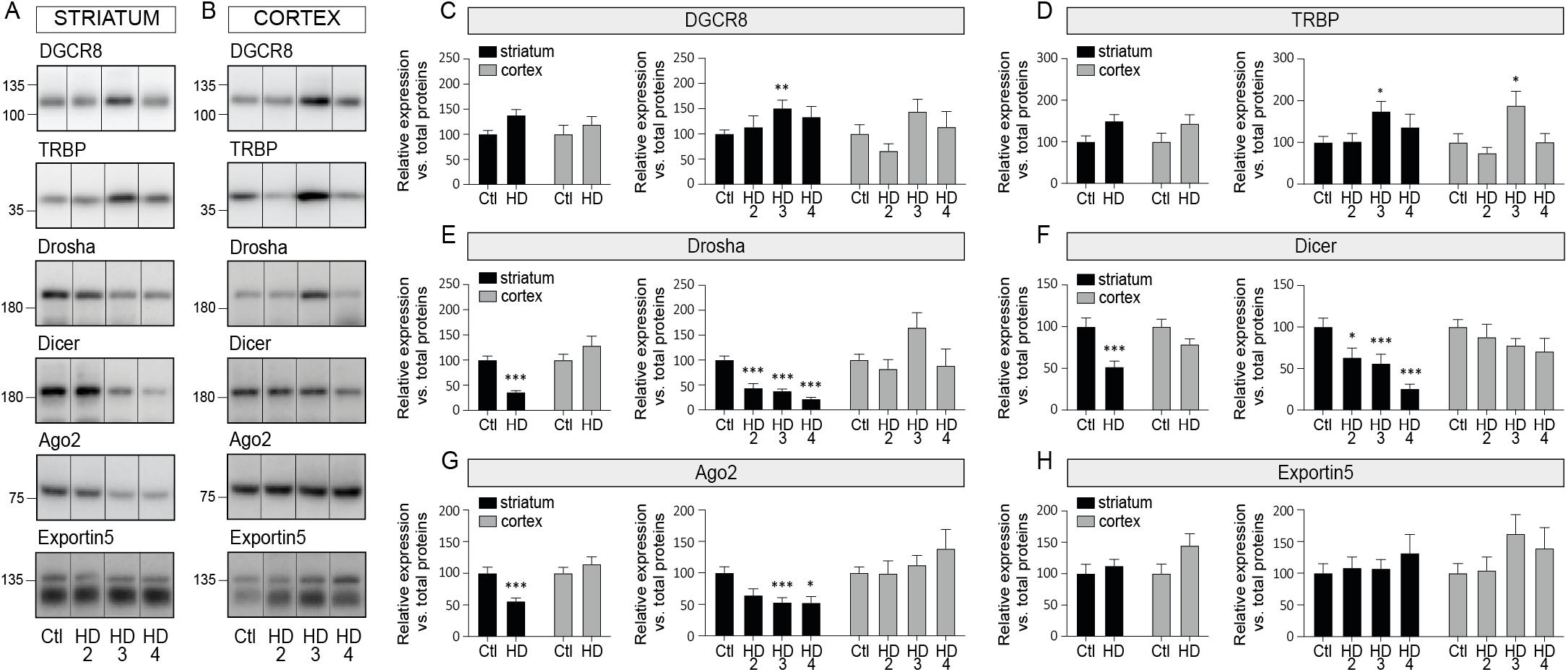
miRNA biogenesis components go awry in human HD striatum. Representative immunoblots of endogenous DGCR8, TRBP, Drosha, Dicer, Ago2 and Exportin in the soluble fraction of **(A)** the striatum or **(B)** the cortex of HD patients and Controls. See Methods for the list of antibodies. **(C-E)** Protein quantifications of each protein according to disease or brain region. Bar graphs with standard error of the mean (SEM) are shown, where the average of Controls is set as 100%. In all cases, the HD group is presented as pooled or per stage. Statistics: Ctl vs. HD as a group was calculated using a Mann-Whitney test. Ctl vs. HD stages was calculated using an analysis of covariance followed by Kruskal-Wallis comparison test. Significant fold changes are provided for each group. * *P*<0.05; ** *P*<0.01; *** *P*<0.001; **** *P*<0.0001. Abbreviations: Ctl, Controls; HD, Huntington’s disease; HD2, Vonsattel grade 2; HD3, Vonsattel grade 3; HD4, Vonsattel grade 4.

### miRNA expression analysis in HD brain

Having shown that several major miRNA biogenesis components were compromised in human HD brain, we next aimed to determine potential effects on miRNA levels. We performed a literature search to identify HD-related miRNAs for downstream functional analyses. Following an initial screen of 16 candidates previously associated with HD, we selected a panel six conserved miRNAs that were commonly misregulated in both HD striatum and cortex (**Fig. 3 and Suppl. Fig. 3**). These included miR-10b, miR-196b, miR-127, miR-128, miR-132 and miR-212. To avoid any bias, we chose miRNAs that were both upregulated and downregulated in HD. Our final choice was further influenced by the different genomic sources of miRNAs: miR-10b and miR-196b are generated from introns of host coding genes, miR-132 and miR-212 are co-expressed as a cluster from the same non-coding gene, miR-127 is expressed from a much larger non-coding miRNA cluster, whereas miR-128 is transcribed from an individual intergenic non-coding gene.

**Figure 3.**
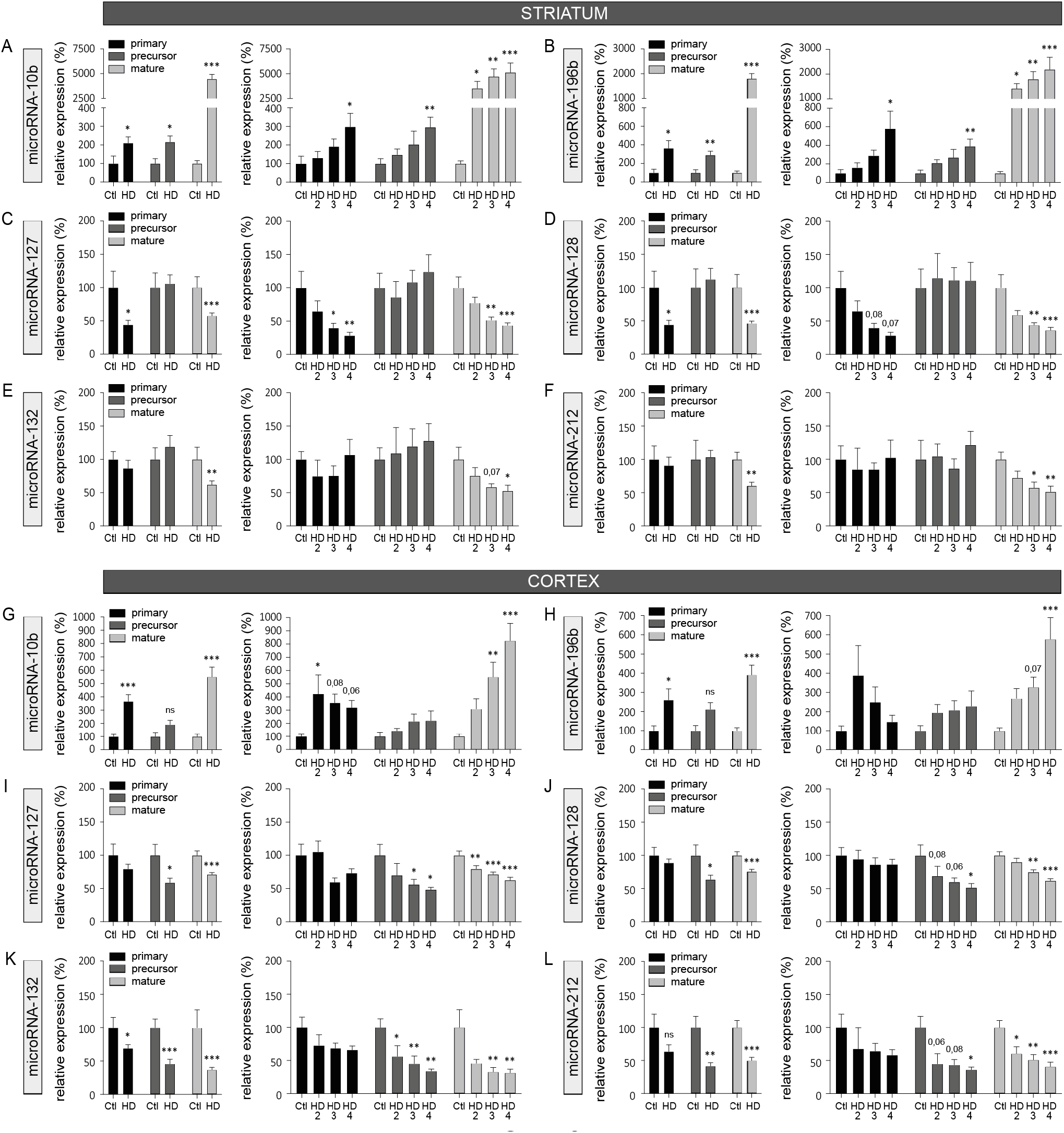
Imbalance between miRNA maturation products in HD brain. **(A-F)** Side-by-side comparison of selected primary, precursor, and mature miRNA transcripts in the striatum of HD patients and Controls. In all assays, we used probe-specific miRNA quantitative RT-PCRs. See Methods for normalization procedures. (**G-L**). A similar analysis of the same miRNAs in matching cortex samples. Bar graphs with standard error of the mean (SEM) are shown, where the average of Controls is set as 100% for each miRNA species. Statistics: Ctl vs. HD as a group was calculated using a Mann-Whitney test. Ctl vs. HD stages was calculated using an analysis of covariance followed by Kruskal-Wallis multiple comparison test. Significant fold changes are provided for each group. * *P*<0.05; ** *P*<0.01; *** *P*<0.001; **** *P*<0.0001. Trends are shown as well. Abbreviations: Ctl, Controls; HD, Huntington’s disease; HD2, Vonsattel grade 2; HD3, Vonsattel grade 3; HD4, Vonsattel grade 4.

We quantified all three types of miRNA maturation products (primary, precursor, mature) in the human striatum and cortex using a distinct set of normalization genes (**Suppl. Fig.4**). As expected, we observed a co-expression of intronic miRNAs and host genes in HD (i.e., miR-10b and miR-19b in the striatum), as documented before (**Fig. 3** and **Suppl. Fig. 2**). Surprisingly, however, various other types of phenomena were observed outside from this canonical pattern, some of which were tissue and disease-stage specific. One example includes the downregulation of pri-miR-127 and mature miR-127, but not pre-miR-127, in HD striatum. Another example includes the specific downregulation of mature miR-132 in HD striatum but an overall downregulation of pri-miR-132, pre-miR-132 and mature miR-132 in HD cortex. In sum, these results suggest that miRNA maturation is controlled at both transcriptional and post-transcriptional levels in HD brain.

### Prominent pre-miRNA to mature miRNA maturation deficits in HD

To better grasp any changes in miRNA maturation *per se* in HD, we analyzed overall ratios (inhibition scores) between a given miRNA precursor and its substrate, as initially proposed by Emde *et al.* [10]. The inhibition scores between pri-miRNA and pre-mRNA were largely normal in HD striatum and cortex despite rare exceptions **(Fig. 4A-F)**. On the other hand, the inhibition scores between pre-mRNA and mature miRNA were drastically altered for all tested miRNAs in HD striatum (**Fig. 4G-L**) whereas only miR-10b and miR-196b reached significance in late-stage HD cortex. Interestingly, miRNA levels and inhibition scores were unaffected in human HD blood in a separate cohort (**Suppl. Fig 5**). Taken together, these results implicate early and robust deficits in pre-miRNA to mature miRNA maturation step in human HD striatum.

**Figure 4.**
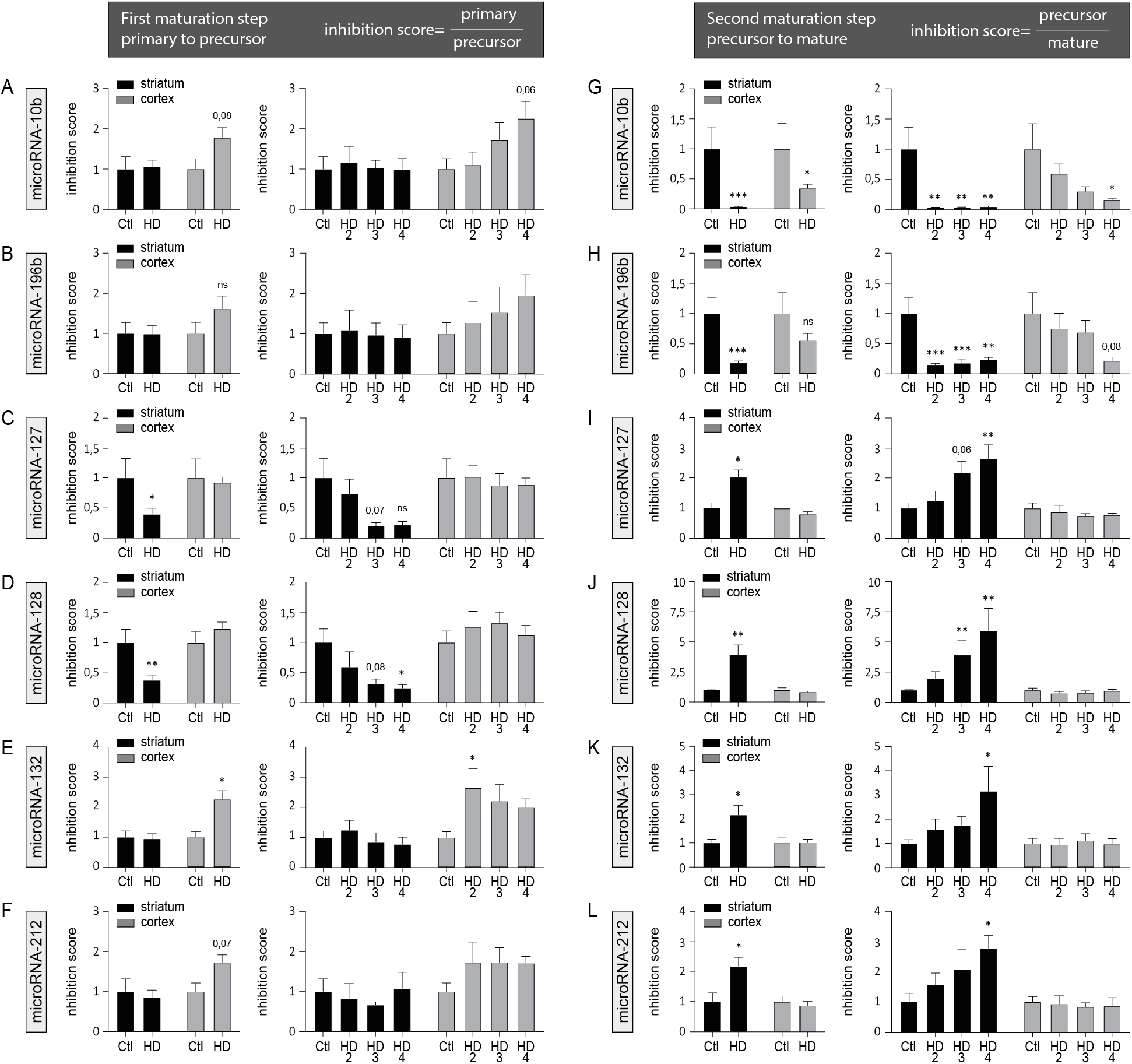
Predominant pre-to mature miRNA maturation deficits in HD brain. **(A-F)** Overview of candidate miRNA primary/precursor inhibition scores (ratios) in the striatum and cortex of HD patients and Controls. Graphs were generated using corresponding qRT-PCR data. Significant differences were observed for a subset of tested miRNAs. **(G-L)** Overview of miRNA precursor/mature inhibition scores (ratios) in the striatum and cortex of HD patients and Controls. Here, all the tested miRNAs were significantly affected, particularly in the striatum. Statistics: Ctl vs. HD as a group was calculated using a Mann-Whitney test. Ctl vs. HD stages was calculated using an analysis of covariance followed by Kruskal-Wallis multiple comparison test. Significant fold changes are provided for each group. * *P*<0.05; ** *P*<0.01; *** *P*<0.001; **** *P*<0.0001. Trends are shown as well. Abbreviations: Ctl, Controls; HD, Huntington’s disease; HD2, Vonsattel grade 2; HD3, Vonsattel grade 3; HD4, Vonsattel grade 4.

**Figure 5.**
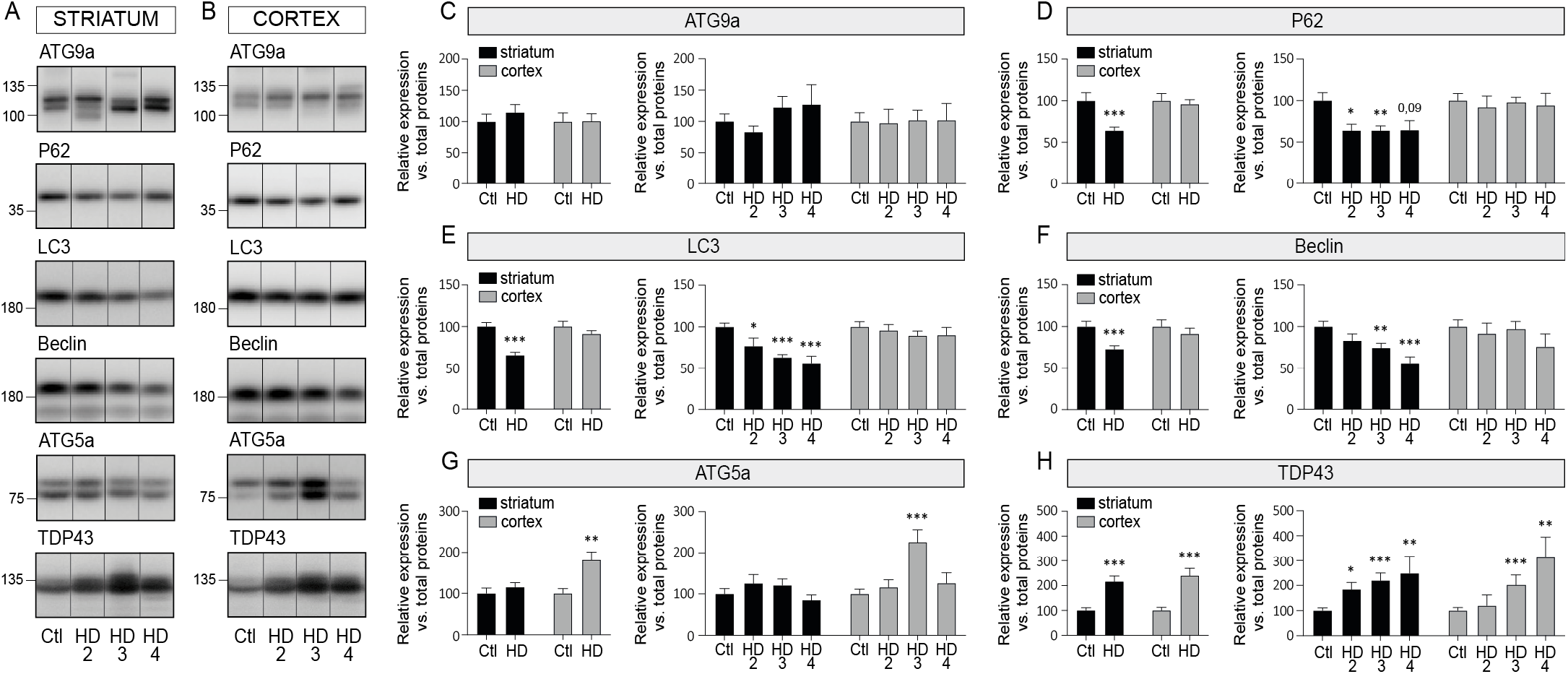
Altered autophagy in HD brain. Representative immunoblots of endogenous ATG9a, P62, LC3, Beclin, ATG5a, and TDP-43 in the soluble fraction of **(A)** the striatum or **(B)** the cortex of HD patients and Controls. See Methods for the list of antibodies. **(C-E)** Protein quantifications of each protein according to disease or brain region. Bar graphs with standard error of the mean (SEM) are shown, where the average of Controls is set as 100%. In all cases, the HD group is presented as pooled or per stage. Statistics: Ctl vs. HD as a group was calculated using a Mann-Whitney test. Ctl vs. HD stages was calculated using an analysis of covariance followed by Kruskal-Wallis multiple comparison test. Significant fold changes are provided for each group. * *P*<0.05; ** *P*<0.01; *** *P*<0.001; **** *P*<0.0001. Abbreviations: Ctl, Controls; HD, Huntington’s disease; HD2, Vonsattel grade 2; HD3, Vonsattel grade 3; HD4, Vonsattel grade 4.

### Autophagy dysfunction overlaps with miRNA maturation defects in HD

Finally, we set out to better understand the molecular mechanisms responsible for miRNA maturation defects in HD. Autophagy dysfunction in an inherent feature of HD and we have previously shown that it influences mature miRNA levels *in vivo* [29]. Accordingly, we observed a strong and significant downregulation of major markers of autophagy, namely P62, LC3 and Beclin, in human HD striatum but not cortex at all stages of the disease (**Fig. 5A-G**). Interestingly, TDP-43 levels, previously implicated in the regulation of miRNA maturation *in vitro* [3, 19], did not correlate with miRNA defects (**5A, B, H**). Taken together, these results strengthen a role for autophagy in modulating miRNA maturation in HD-susceptible brain regions.

## Discussion

This study provides the first *in vivo* evidence that miRNA maturation is dysregulated in human HD and sheds a new light on the causes and potential implications of miRNA dysregulation in HD. The importance of our findings is severalfold: 1) they provide important clues on intra-individual variability and susceptibility towards mHtt pathology, 2) they could explain a substantial proportion of miRNA alterations previously documented in HD brain, 3) they provide a first in-depth analysis of the RISC components necessary for endogenous RNA interference (RNAi), 4) they support the potential importance of specific miRNAs (and downstream targets) in HD pathogenesis, and finally 5) they strengthen the broad implications of autophagy dysregulation in HD pathogenesis.

To our knowledge, this is the first characterisation of human Htt expression and aggregation in two different brain regions affected in HD from the same individuals. These experiments validate and extend our previous biochemical studies focused on Htt pathology and other proteinopathies exclusively in the striatum (putamen) [40]. In agreement with earlier reports, lower Htt (mHtt) expression (loss-of-function) and higher mHtt aggregation (gain-of-function) are likely both contributing factors to the severe neurodegeneration observed in the striatum. It is interesting to note that (at least some of) the proposed toxic Htt N-terminal fragments [31, 44] were upregulated in both brain regions analyzed, suggesting that additional or complementary factors participate in Htt-mediated toxicity. This hypothesis is consistent with a role for miRNA-dependent survival pathways in this process.

Remarkably, very little is known about the underlying causes of miRNA alterations in HD, which is key to understanding the role, impact, diagnostic, and therapeutic potential of miRNAs in human brain diseases. Clearly, the inhibitory effects of mHtt on transcription [21, 30, 33] are readily evident in this study on both coding (e.g., miR-10b) and non-coding (e.g., miR-127) genes their host miRNA transcripts. In addition to transcriptional effects, our observations implicate other molecular mechanisms as major causes of mature miRNA disruption in HD. The identification of factors that control pre-miRNA to miRNA maturation abnormalities in HD (e.g., transport, cleavage, sequestration, degradation) will require further investigation. Interestingly, cellular stress has been shown to disrupt pre-miRNA to mature miRNA genesis in ALS [10]. Stress can influence miRNA maturation in several ways, including the sequestration of pre-miRNAs and pathway components (e.g., Ago2) to P bodies and/or stress granules. On this line of thought, autophagy is functionally implicated in mHtt protein turnover and aggregation, and more recently miRNA maturation [2, 43]. Without a doubt, more studies are required to understand the cause-and-effect relationship between these factors during HD progression.

Interestingly, most miRNA biogenesis components were downregulated in human HD striatum. This observation is somewhat consistent with earlier findings in mice that showed a transient shift (up to down) in miRNA expression levels during disease progression. It remains to be elucidated whether the triggering factor is a unique component (e.g., Ago2 downregulation [28]) or a more general mechanism in humans. The study of pre-symptomatic HD patients (i.e., Vonsattel grades HD0-1) or humanized cell models (e.g., iPSC) would undoubtedly help to address this question. In any case, our results are consistent with an abnormal regulation of miRNA biogenesis in HD.

Of note, we did not observe changes in mature miRNA levels (not shown) or miRNA maturation defects in HD blood, although we and others have previously reported high expression levels of Htt in blood cells [7, 32], further strengthening the hypothesis of tissue-specific effects. At this stage, however, we cannot exclude maturation defects for other miRNAs and/or cohorts. A critical question is how do mature miRNAs become misregulated in tissues or cell types with seemingly normal miRNA biogenesis. As shown herein, changes in gene transcription can lead to alternations of miRNA host genes and henceforth mature miRNA output. In addition, and as mentioned above, mature miRNA levels are subjected to multiple regulatory mechanisms (e.g., degradation) and feedback loops that can go awry in disease conditions as well. An interesting hypothesis is that the specific disruption of miRNA biogenesis – and not indirect effects of neurodegeneration on mature miRNA levels – is responsible for the early susceptibility of cell loss in HD. This could have context-specific consequences on key miRNAs or other RISC-dependent RNAs that are required to maintain cell homeostasis.

On this line of thought, several groups have already tested the regulatory effects of candidate miRNAs on HD pathology, behavior, and cell survival. For example, an increased expression of miR-196a (homologue of 196b) in transgenic mice caused lower mHtt expression and aggregation in an HD model [4]. Overexpression of miR-10b in PC12 cells expressing mHtt also increased cell survival [16]. Finally, the brain supplementation of miR-132 in HD mice partially rescued behavioral and motor symptoms [11]. Interestingly, the miR-132/212 cluster is among the most strongly affected miRNA (family) in HD brain (this study and [11, 22]). We have previously shown that miR-132/212 knockout mice display autophagy abnormalities and lower BDNF levels in the brain as seen in HD [14, 15, 34, 45]. Additional studies are also required to establish the underlying causes of Drosha, Dicer and Ago2 downregulation in HD striatum, although autophagy is a reasonable candidate as well. The challenge now is to identify the targets and pathways regulated by mature and possibly immature miRNA transcripts for in-depth functional analyses *in vivo*, of course taking into account the occurrence of potential transient changes as observed in HD mice and tissue-specific effects.

Interestingly, recent evidence suggests that impaired miRNA maturation occurs in other trinucleotide repeat disorders as well. For instance, the expanded CGG repeats in FMRP (causing Fragile X-associated tremor/ataxia syndrome) sequester DGCR8 and Drosha and disrupt miRNA maturation in mice [38]. In drosophila, mutant ataxin-2 (causing spinocerebellar ataxia type 2) disrupts Ago expression and miRNA function [26]. MiRNA maturation is impaired also in models of expanded polyQ within ataxin-3 (causing Machado-Joseph disease), whereas blocking miRNA biogenesis increased ataxin-3 aggregation [20]. These observations strongly suggest that miRNA alterations in these disorders are a direct consequence of disease genes (e.g., sequestration) rather than an indirect effect of cell stress or other factors. The fact that Htt binds to Ago2 is consistent with this hypothesis although a role for additional genetic or molecular mechanisms cannot be excluded in these diseases.

The endogenous RISC complex is central to the silencing of genes by miRNAs and other small interfering RNAs such as small interfering RNAs (siRNAs). Interestingly, a variety of therapeutic tools under development make use of miRNAs, siRNAs, or other antisense oligonucleotides (including miRNA-like backbones) that silence mHtt expression *in vivo* [1, 8]. As shown herein, the clinical testing of these compounds in human brain will need to be carefully monitored for potential loss of RISC biological function. Clearly, much more work is required to better understand the role and impact of miRNA biogenesis abnormalities in HD and related trinucleotide disorders.

## Conclusions

In summary, we show that pre-miRNA to mature miRNA biogenesis is strongly compromised in human HD striatum. This observation could help to understand the pathological relationship between Htt-Ago2 binding *in vivo*. Furthermore, this study suggests that indirect or small changes in mature miRNA levels are not sufficient to promote cell degeneration *per se* in trinucleotide diseases, compared to a “multiple-hit” scenario implicating deficits in miRNA biogenesis or other RISC-dependent mechanisms. Finally, the results provided herein will guide current and future therapeutic strategies involving the endogenous RISC in the human brain.

## List of abbreviations

DGCR8: DiGeorge syndrome critical region 8
TRBP: Human immunodeficiency virus transactivating response RNA-binding protein
Ago2: Argonaut 2
RISC: RNA-induced silencing complex
NeuN: Neuronal nuclear protein
DARPP-32: Dopamine- and cAMP-regulated neuronal phosphoprotein
PSD95: Postsynaptic density protein 95
BDNF: Brain-derived neurotrophic factor
nt: nucleotide

## Acknowledgement

This work was supported by the Canadian Institute of Health Research (CIHR, grant # 272311), the Fonds de Recherche du Québec en Santé (FRQS), and the Huntington’s Disease Society of America (HDSA). The Harvard Brain Tissue Resource Center provided tissues and is supported in part by HHSN-271-2013-00030C. The authors would like to express a special appreciation to the nurses and staff who assisted in the collection and storage of the human specimens. Finally, the authors are very grateful to all study participants and their families who have contributed to this study.

## Ethics approval and consent to participate

The experiments with human samples were approved by the CHU de Québec local Research Ethics Committee (#2017-3017). All work with human subjects was approved by the CHU de Québec human ethics committee (#2020-4622) and in accordance with the Declaration of Helsinki. Informed written consent was obtained from all participants.

