## Supplementary Material for "Widespread alterations in microRNA biogenesis in human Huntington’s disease putamen"

### Supplementary Methods

#### Human blood samples

All work with human subjects was approved by the CHU de Québec human ethics committee (#2020-4622) and in accordance with the Declaration of Helsinki. This cohort was recruited from the Clinique des troubles du mouvement of the CHU de Québec, Québec City, Canada. Informed written consent was obtained from all participants. A total quantity of 4 ml of blood was collected from patients with HD at all stages of disease (n=8) along with healthy control subjects (n=7) for a total of 15 participants (Supplemental Table 2). Clinical evaluations were conducted on the same day of blood sampling. Whole blood: Total RNA was isolated from 500 µl of whole blood using the TRIzol reagent (Ambion by Life Technologies) according to the manufacturer's instructions. PBMCs: peripheral blood mononuclear cells (PBMCs) were first isolated from 3 ml of whole blood using Ficoll-Plaque PLUS (GE Healthcare, 71-7167-00 AG) according to manufacturer's instruction. Then, purification of total RNA was extracted from PBMCs using TRIzol reagent (Ambion by Life Technologies) according to the manufacturer's instructions. RNA was then stored at -80°C until use.

**Supplementary Tables**

**Table 1. Characteristics of the individuals providing post-mortem brain samples (NIH Biobank)**

| | N | Age | PMI | Women (%) | Men (%) | Allele 1 | Allele 2 | Disease score |
|---|---|---|---|---|---|---|---|---|
| Control | 25 | 67 [35-79] | 20 [8-24] | 40 | 60 | 21 | 24 | - |
| HD2 | 10 | 64 [49-80] | 21 [12-27] | 60 | 40 | 22 | 47 | 703 |
| HD3 | 23 | 59 [47-75] | 20 [8-27] | 39 | 61 | 25 | 48 | 726 |
| HD4 | 8 | 52 [47-64] | 19 [12-24] | 50 | 50 | 23 | 52 | 844 |
| HD | 41 | 59 [43-75) | 20 [12-27] | 46 | 54 | 24 | 49 | 758 |

**Table 3. Antibodies used in this study.**

| | Antibody | Provider | Cat number | Species | Dilution | Proteins | System |
|---|---|---|---|---|---|---|---|
| Huntingtin | Anti-Htt (CHU00146) | Coriell Institute | CHDI-90000137 | Rabbit | 1/1000 | 20 µg | Tris-acetate 3-15% |
| Htt total | Anti-Htt (1HU-4C8) | Millipore | mab2166 | Mouse | 1/1000 | - | Dot blot |
| Htt poly-Q | Anti-polyglutamine expansion (5TF1-1C2) | Millipore | mab1574 | Mouse | 1/1000 | - | Dot blot |
| NeuN | Anti-NeuN (E4M5P) | Cell Signaling Technology | 94403 | Mouse | 1/10 000 | 5 µg | 10% acrylamide |
| Darpp32 | Anti-DARPP-32 (19A3) | Cell Signaling Technology | 2306 | Rabbit | 1/10 000 | 5 µg | 10% acrylamide |
| PSD95 | Anti-PSD95 | Cell Signaling Technology | 2507 | Rabbit | 1/1000 | 10 µg | 10% acrylamide |
| DGCR8 | Anti-DGCR8 [EPR18757] | Abcam | ab191875 | Rabbit | 1/1000 | 10 µg | 10% acrylamide |
| TRBP | Anti-TRBP [EPR13550] | Abcam | ab180947 | Rabbit | 1/1000 | 10 µg | 10% acrylamide |
| Exportin5 | Anti-Exportin 5 (D7W6W) | Cell Signaling Technology | 12565 | Rabbit | 1/1000 | 10 µg | 10% acrylamide |
| Dicer | Anti-Dicer (F-10) | Santa Cruz Biotechnology | sc-136979 | Mouse | 1/1000 | 10 µg | 10% acrylamide |
| Drosha | Anti-Drosha (D28B1) | Cell Signaling Technology | 3364 | Rabbit | 1/1000 | 10 µg | 10% acrylamide |
| Ago2 | Anti-Argonaute 2 (C34C6) | Cell Signaling Technology | 2897 | Rabbit | 1/1000 | 10 µg | 10% acrylamide |
| ATG9a | Anti-Atg9a (D4O9D) | Cell Signaling Technology | 13509 | Rabbit | 1/1000 | 10 µg | 10% acrylamide |
| P62 | Anti-SQSTM1/p62 | Cell Signaling Technology | 5114 | Rabbit | 1/1000 | 10 µg | 10% acrylamide |
| LC3 | Anti-LC3B | Novus Biological | NB 100-2220 | Rabbit | 1/1000 | 20 µg | Tris-acetate 3-15% |
| Beclin | Anti-Autophagy pack | Novus Biological | NB 910-94877 | Rabbit | 1/1000 | 10 µg | 10% acrylamide |
| ATG5 | Anti-ATG5 | Novus Biological | NB 110-53818 | Rabbit | 1/1000 | 10 µg | 10% acrylamide |
| TDP43 | Anti-Tdp43 (3H8) | Millipore | mabN45 | Rabbit | 1/1000 | 10 µg | 10% acrylamide |

### Supplementary Figures

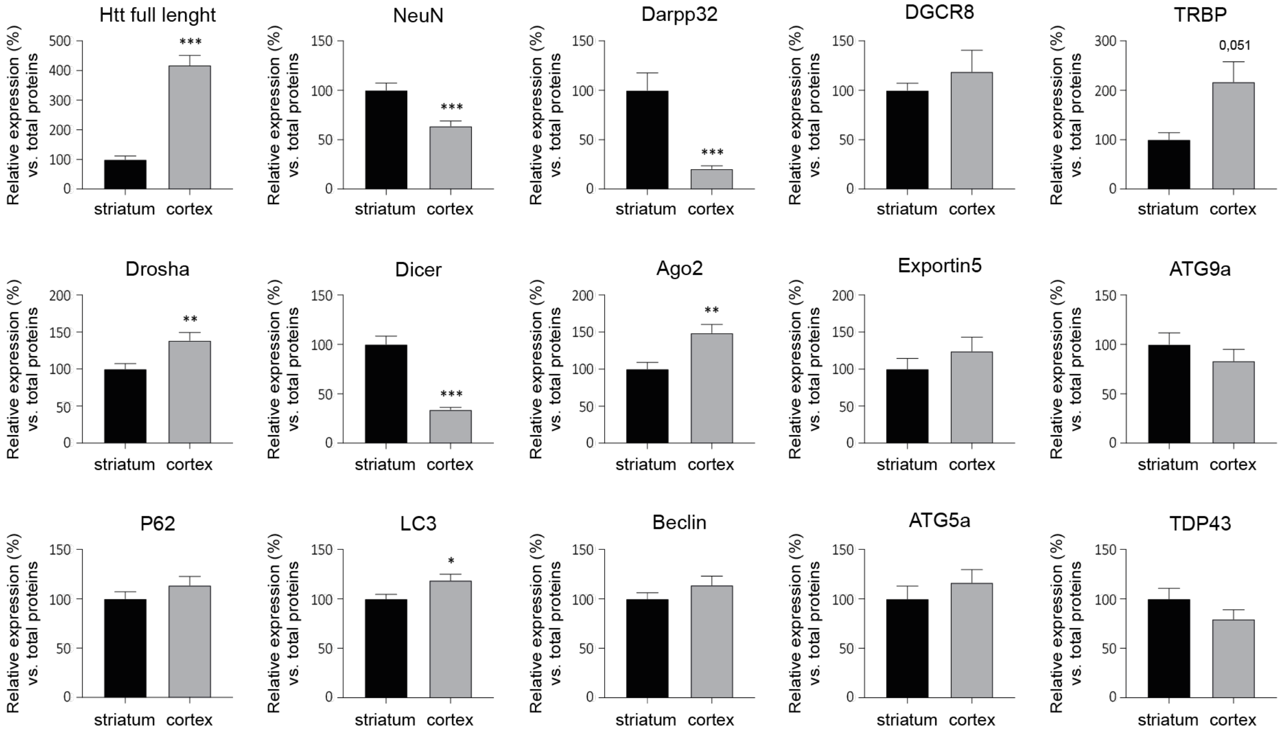


**Figure 1. Comparative analysis of protein expression between brain regions.** Shown here are Western blot quantifications of different proteins tested in this study. Soluble proteins extracted from the striatum and the cortex (N=25 healthy controls) were loaded onto the same gel for a side-by-side comparison. Bar graphs with standard error of the mean (SEM) are shown, where the average of the striatum is set as 100%. Statistics: Striatum vs. Cortex was calculated using a Mann-Whitney test. * $P<0.05$; ** $P<0.01$; *** $P<0.001$; **** $P<0.0001$. Trends are shown as well.

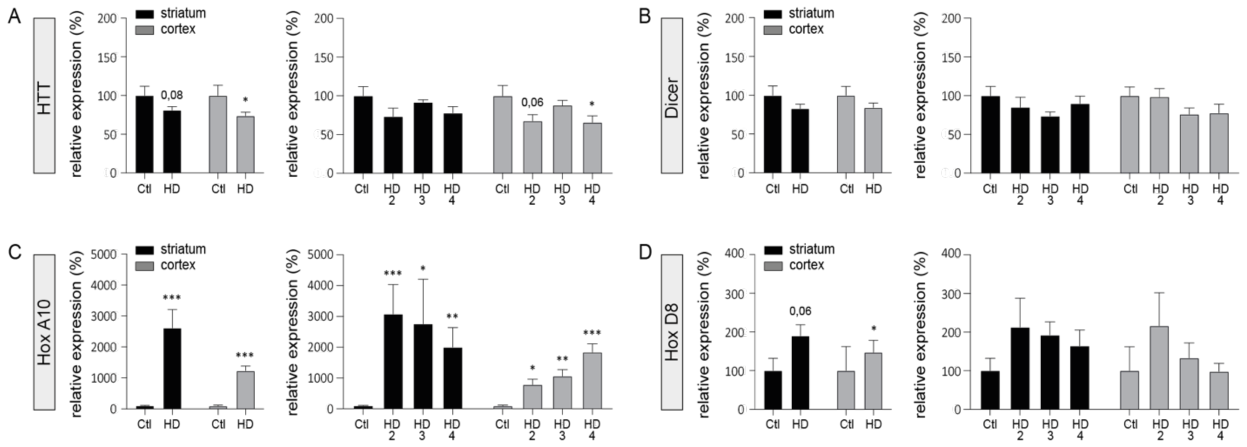


**Figure 2. Analysis of mRNA expression in HD brain.** qRT-PCR analysis of endogenous Htt, Dicer, HoxA10 and HoxD8 mRNA in HD patients (HD2 N=9; HD3 N=9 and HD4 N=8)) and healthy controls (N=9). Statistics: Ctl vs. HD as a group was calculated using a Mann-Whitney test. Ctl vs. HD stages was calculated using an analysis of covariance followed by Kruskal-Wallis multiple comparison test. Significant fold changes are provided for each group. * $P<0.05$; ** $P<0.01$; *** $P<0.001$; **** $P<0.0001$. Trends are shown as well. Abbreviations: Ctl, Controls; HD, Huntington's disease; HD2, Vonsattel grade 2; HD3, Vonsattel grade 3; HD4, Vonsattel grade 4.

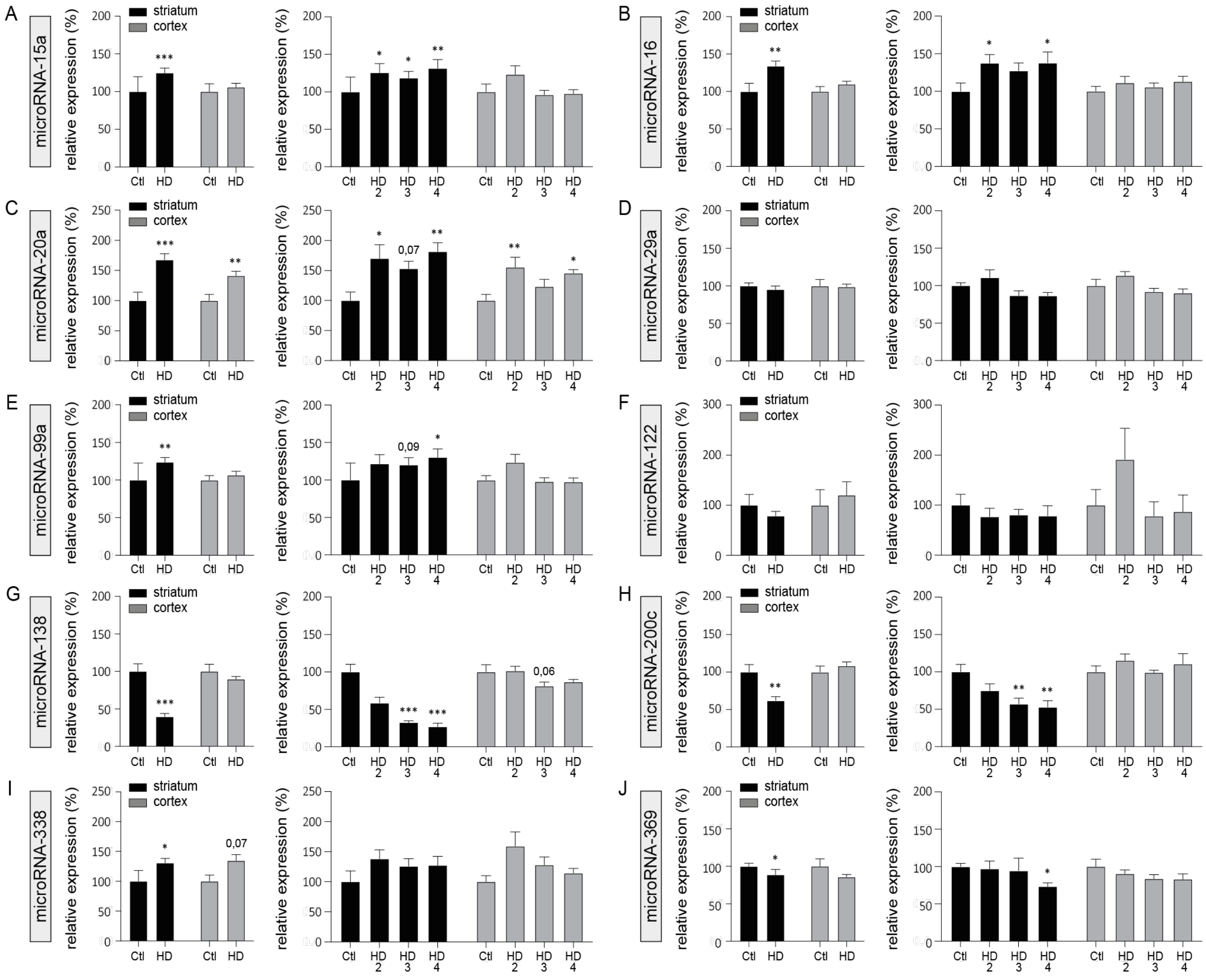


**Figure 3. miRNA screening in HD brain.** The relative expression levels of additional HD-related mature miRNAs were evaluated by miRNA qRT-PCR in HD patients (HD2 N=9; HD3 N=9 and HD4 N=8) and healthy controls (N=9). Most changes occurred in the striatum. Statistics: Ctl vs. HD as a group was calculated using a Mann-Whitney test. Ctl vs. HD stages was calculated using an analysis of covariance followed by Kruskal-Wallis multiple comparison test. Significant fold changes are provided for each group. * $P<0.05$; ** $P<0.01$; *** $P<0.001$; **** $P<0.0001$. Trends are shown as well. Abbreviations: Ctl, Controls; HD, Huntington's disease; HD2, Vonsattel grade 2; HD3, Vonsattel grade 3; HD4, Vonsattel grade 4.

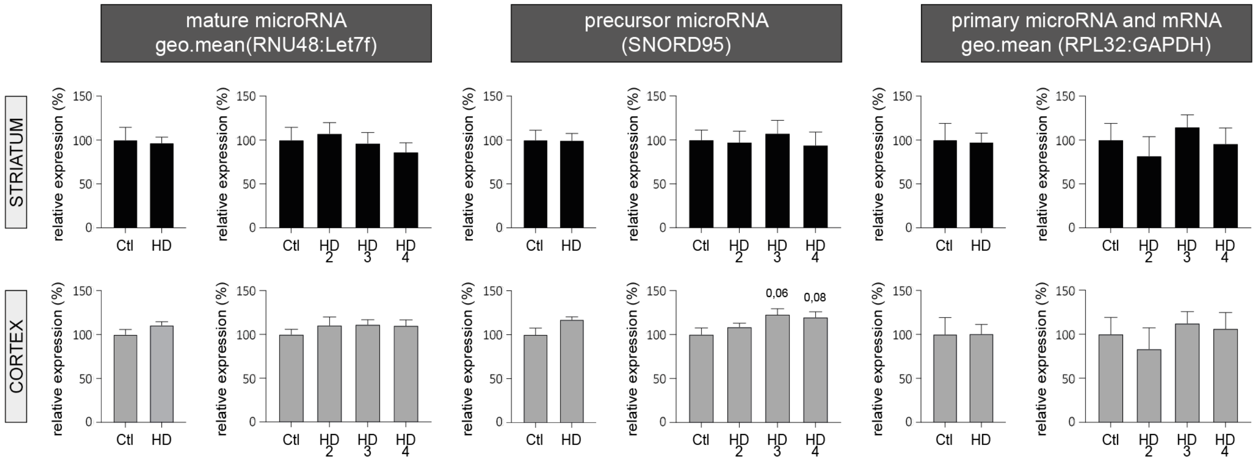


**Figure 4. qRT-PCR analysis of normalization genes used in this study.** Quantifications were done using both HD patients (HD2 N=9; HD3 N=9 and HD4 N=8) and healthy controls (N=9). Statistics: Ctl vs. HD as a group was calculated using a Mann-Whitney test. Ctl vs. HD stages was calculated using an analysis of covariance followed by Kruskal-Wallis multiple comparison test. No statistically significant differences were observed overall. Abbreviations: Ctl, Controls; HD, Huntington's disease; HD2, Vonsattel grade 2; HD3, Vonsattel grade 3; HD4, Vonsattel grade 4.

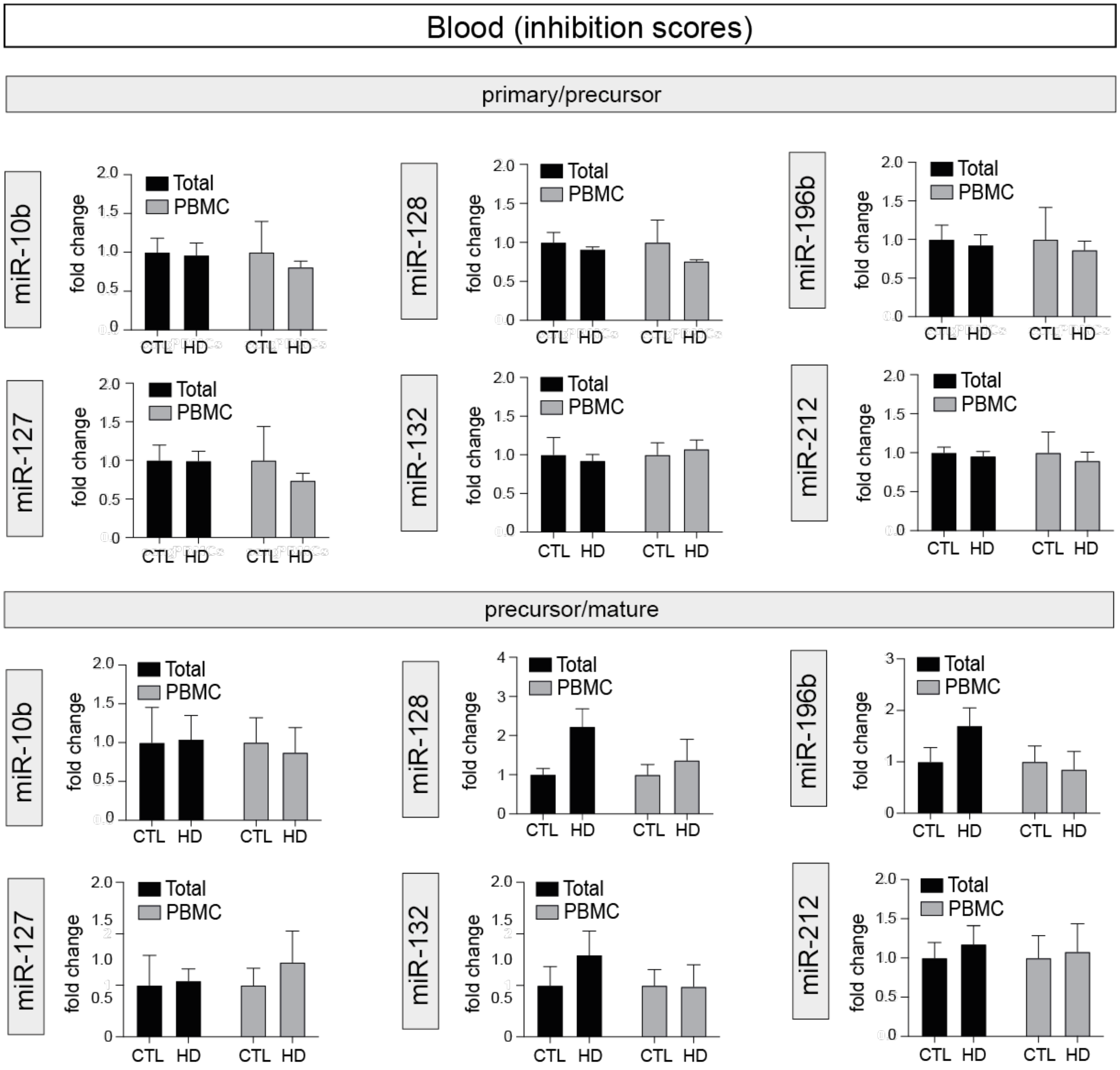


**Figure 5. Analysis of miRNA maturation in blood.** Overview of miRNA inhibition scores in human blood of HD (N=7) and healthy control (N=8) subjects of the CHU de Québec cohort. Here, we used total RNA extracted from either whole blood or PBMCs. No significant effects were observed for all tested miRNAs. In addition, no changes in mature miRNA levels were noted in these samples (not shown). Statistics: Ctl vs. HD as a group was calculated using an ANOVA test with multiple comparisons.
